# A long-non-coding RNA, *LINC00473*, confers the human adipose tissue thermogenic phenotype through enhanced cAMP responsiveness

**DOI:** 10.1101/339192

**Authors:** Khanh-Van Tran, Cecilie Nandrup-Bus, Tiffany DeSouza, Ricardo Soares, Naja Zenius Jespersen, So Yun Min, Raziel Rojas-Rodriguez, Hanni Willenbrock, Therese Juhlin, Mai Charlotte Krogh Severinsen, Kimberly Malka, Bente Klarlund Pedersen, Timothy Fitzgibbons, Camilla Scheele, Silvia Corvera, Søren Nielsen

## Abstract

Specialized adipocytes localized in distinct depots mediate the many physiological functions of adipose tissue. In humans, paucity of thermogenic adipocytes correlates with high metabolic disease risk, raising much interest in the mechanisms by which these cells arise. Here we report molecular signatures associated with adipocyte development in different human depots and identify a long non-coding RNA, *LINC00473*, as the transcript most closely associated with enrichment of thermogenic adipocytes. *LINC00473* expression is low in subjects with obesity or type-2 diabetes and is highly correlated with cAMP signaling and mitochondrial oxidative phosphorylation pathways. *LINC00473* is localized in the nucleus and the cytoplasm, and its knockdown impairs induction of UCP1 and mitochondrial respiration. These results reveal that depot-enriched genes that modulate responsiveness to external stimuli, specifically *LINC00473*, are important determinants of the adipose tissue thermogenic phenotype, and potential targets for metabolic disease therapy.

## Introduction

Adipose tissue is a central player in the control of whole body energy homeostasis, playing numerous roles including energy storage and release, endocrine control of fuel homeostasis and thermogenesis (Crewe et al., 2017; Harms and Seale, 2013; Nedergaard and Cannon, 2010; Rosen and Spiegelman, 2006). These functions are mediated by specific adipocyte subtypes, which are enriched in specific adipose tissue depots. For example, the majority of thermogenic, Ucp1 containing adipocytes in mice are localized in a specific depot within the interscapular region, which is mostly devoid of white adipocytes (Cinti, 2001). In adult humans, thermogenic adipocytes are interspersed amongst ostensibly white, non-thermogenic cells forming a “brown-in-white” (brite) or “beige” depot, which is found in the supraclavicular and paravertebral region (Cypess et al., 2013; Jespersen et al., 2013; Virtanen et al., 2009). Thus, both in mouse and human, specialization of adipocyte function is associated with defined anatomical localization.

The anatomical localization of adipocytes is relevant for adipose tissue development, as shown by findings that adipocyte progenitors transplanted into different regions of mice give rise to functionally different adipocytes (Jeffery et al., 2016). In addition, adipocytes within different depots descend from distinct embryonic mesenchymal precursor cells (Sanchez-Gurmaches et al., 2016), and this lineage can specify distinct pools of brown and white adipocytes in different depots. The molecular mechanisms dictated by the combination of lineage determinants and local anatomical environments are not fully understood, but are likely to include transcriptional regulatory mechanisms that have been associated with development of different adipocyte phenotypes (Mota de Sa et al., 2017).

Recent studies have brought attention to the role of long non-coding RNAs in the establishment of adipocyte functions (Alvarez-Dominguez et al., 2015; Bai et al., 2017; Zhao et al., 2014). Long noncoding RNAs are endogenous cellular RNAs of more than 200 nucleotides in length that lack open reading frames. The long non-coding RNAs located as an independent unit between two coding genes are termed long intergenic non-coding RNA (Linc RNA) (Atianand et al., 2017; Freedman and Miano, 2016; Gutschner and Diederichs 2012). Recent transcriptome analyses have revealed thousands of non-coding RNAs which can potentially regulate gene expression at multiple levels, including chromatin modification, transcription and post transcriptional processing (Faghihi et al., 2008; Gupta et al., 2010; M et al., 2010; Mercer et al., 2009; Simon et al., 2011; Tsai et al., 2010; Wang et al., 2008). In mice, a nuclear lncRNA (Blinc1) was implicated in the development of brown and beige adipocytes through a ribonucleoprotein complex containing the transcription factor EBF2 (Zhao et al., 2014). Another lncRNA, lncBATE10, can prevent repression of Pgc1alpha mRNA and sustain the thermogenic phenotype (Alvarez-Dominguez et al., 2015; Bai et al., 2017). Thus, long nocoding RNAs could be central players in integrating anatomical and lineage factors to produce functionally distinct adipocytes residing in different depots.

The extent to which mechanisms of adipocyte development are conserved between species is not known. In particular, regulatory mechanisms dependent on non-coding RNAs may vary, as there is limited conservation and large plasticity between species (Hezroni et al., 2015). This information is relevant in the context of metabolic disease, since increased metabolic activity of thermogenic adipose tissue has been linked with leanness, increased energy expenditure and improved glucose homeostasis in humans (Scheele and Nielsen, 2017). Understanding the mechanisms that determine the anatomical localization and abundance of thermogenic adipocytes could reveal pathogenic underpinnings of metabolic disease and suggest therapeutic targets. In mice, the ability to perform genetic lineage tracing has been harnessed for identification of adipocyte progenitors and their fate (Berry and Rodeheffer, 2013; Sanchez-Gurmaches et al., 2016; Tran et al., 2012). While lineage tracing cannot be performed in humans, relevant information can be derived from the analysis of multipotent mesenchymal progenitors that are present within the tissue, and give rise to new adipocytes. In previous studies, we observed that human adipocyte progenitor cells are associated with the adipose tissue microvasculature, and proliferate robustly in response to angiogenic stimuli (Min et al., 2016). Furthermore, brown fat differentiation program is cell autonomous and largely depot-dependent (Jespersen et al., 2013). We now leverage these finding to determine whether progenitors and the adipocytes they give rise to from different adipose depots display distinct genetic signatures, and which signatures are specifically associated with the development of the thermogenic phenotype.

## Results

To elucidate the mechanisms involved in the generation of thermogenic adipocytes in humans, we first searched for major gene expression differences between adipocytes generated from non-thermogenic and thermogenic adipose tissue depots. Biopsies from periumbilical (AbdSQ) and from supraclavicular (SClav) adipose tissue of non-diabetic subjects were obtained (Supplementary table 1, subject characteristics). Cells were extracted by collagenase digestion, differentiated as depicted in Figure 1a, and stimulated for 4h with norepinephrine (NE) prior to RNA extraction. Differentiation was seen in ~80% of cells, as assessed by accumulation of lipid droplets (Figure 1b). RNASeq of SClav and AbdSQ derived cells identified 29,907 annotated genes, which segregated in the first principal component into two main groups corresponding to the two depots of origin (Figure 1c). Unsupervised hierarchical clustering of the top 1000 most varied genes also produced two main clusters (Figure 1d), with further segregation within each cluster. Segregation within the AbdSQ cluster was determined by individual subjects, while segregation within the SClav cluster was determined by NE stimulation. Thus, large NE-induced gene expression changes in SClav mitigated subject-defined differences. Nevertheless, we identified a group of genes that responded to NE in cells from both AbdSQ and SClav from all subjects. This group of genes contained the long non-coding RNA, *LINC00473* (Figure 1d). Differential expression analysis identified *LINC00473* to be among the most strongly induced of the 313 genes significantly regulated by NE (Figure 1e,f and Supplementary table 2). For comparison, the levels of genes that have been previously associated with brown or white adipose tissue are shown (Figure 1f). As expected, *HOXC8* and *HOXC9* were expressed at significantly higher levels in AbdSQ than SClav, and *TBX1* displayed the opposite trends. *UCP1* and *DIO2* were detected in SClav prior to NE stimulation, consistent with the presence of ‘brown’ adipocytes in this compartment, as defined by the presence of *UCP1* prior to adrenergic stimulation.

**Figure 1.**
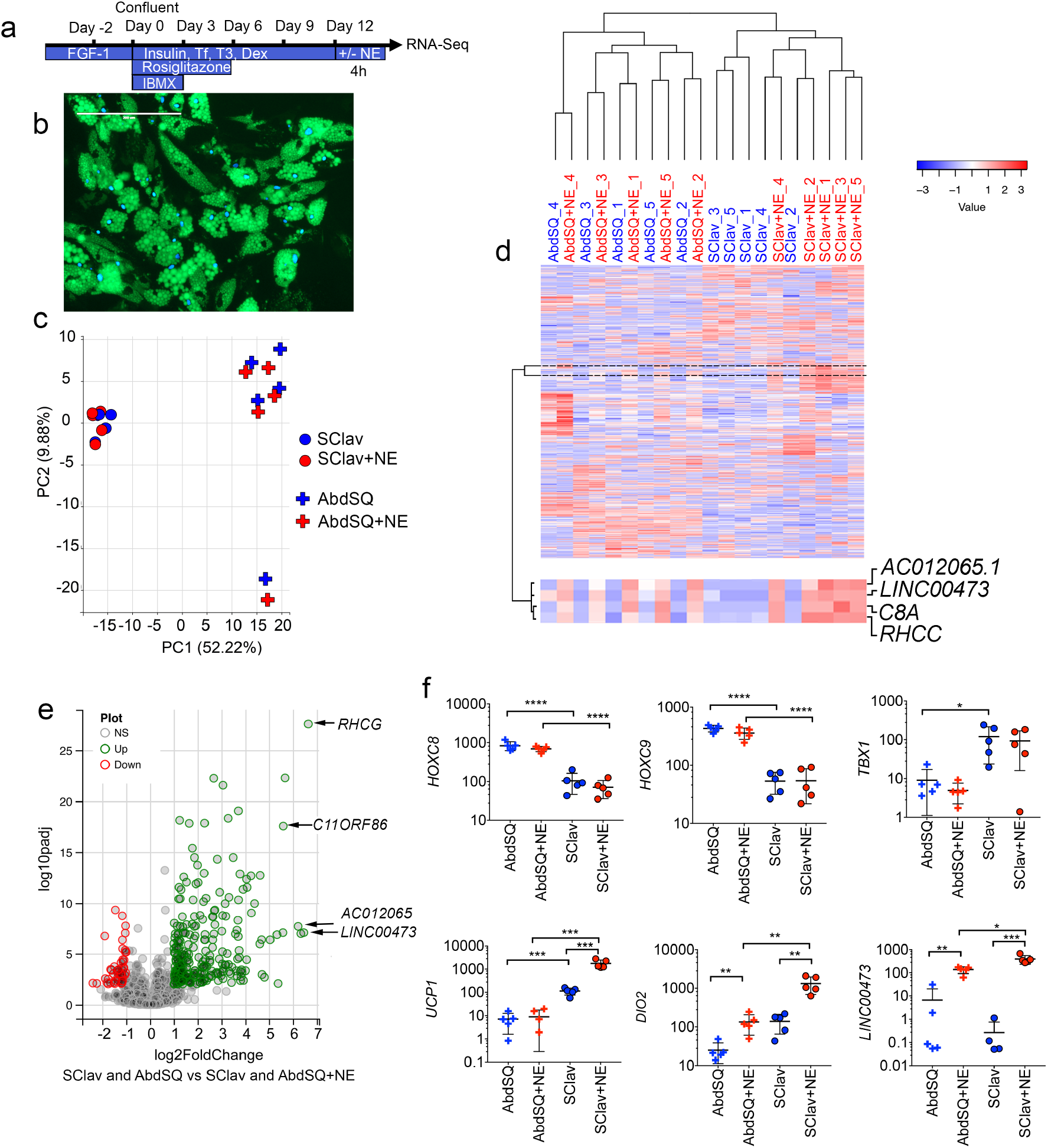
Norepinephrine effects on gene expression in primary adipocytes from thermogenic and non-thermogenic human adipose tissue. **a**. Scheme for differentiation of primary adipocytes from stromovascular fraction. **b**. Example of differentiated adipocytes from AbdSQ depot, showing lipid roples marked by Bodipy (green), and nuclei labeled with Dapi (blue). **c**. Principal component analysis of the 1000 most differentially expressed genes between depots obtained by RNAseq of primary adipocytes from supraclavicular (SClav) or abdominal subcutaneous (AbdSQ) adipose tissue from 5 independent subjects, without or with treatment with norepinephrine (NE). **d**. Unsupervised hierarchical clustering using Pearson’s correlation of same gene set. Marked is a cluster of genes exhibiting increases expression in response to NE in all depots. **e**. Volcano plot of comparison between minus and plus NE in both depots. **f**. FPKM values for selected genes. Plotted are means and SEM. Statistical differences were calculated using multiple t-tests, corrected for multiple comparisons using the Holm-Sidak method. *=p<0.05; **p=<0.01; ***p<0.001; ****p< 0.0001.

To probe for genes involved in the development of thermogenic adipose tissue, we leveraged our previous finding that progenitor cells within the vasculature of AbdSQ adipose tissue robustly proliferate in 3-dimensional culture under pro-angiogenic conditions. We first tested whether progenitors could be generated from subcutaneous adipose tissue from the neck (NeckSQ), which is enriched in thermogenic adipocytes (Cypess et al., 2013), and from perivascular adipose tissue obtained from the carotid artery (Carotid). We find that after approximately 5 days in culture, explants from these depots generated sprouts, which grew robustly for a minimum of 14 days (Figure 2a). We then proteolytically dissociated the cultures, grew cells to confluence and exposed them to a minimal adipogenic cocktail (MDI: methylisobutyl xanthine, dexamethasone and insulin). Adipocyte progenitors readily differentiated into adipocytes as assessed by the accumulation of large lipid droplets (Figure 2b, middle panels), which reduced in size in response to Forskolin (Fsk) stimulation (Figure 2b, right panels). The adipocyte-specific genes *PLIN1, ADIPOQ* and *FABP4* were similarly induced upon differentiation of progenitors from both depots (Figure 2c-e). In contrast, *UCP1* was induced upon differentiation of NeckSQ progenitors, but not of Carotid perivascular progenitors (Figure 2f). These results indicate that progenitors for ‘brown’ adipocytes, as defined by the expression of *UCP1* prior to thermogenic stimulation, are present at the highest level in the NeckSQ depot in our cohort. Nevertheless, progenitors for inducible “brite/beige” adipocytes were present in both depots, as evidenced by strong expression of *UCP1* in response to Fsk stimulation of adipocytes differentiated from Carotid or NeckSQ progenitors (Figure 2g). The relationship between the numbers of ‘brown’ and ‘beige/brite’ progenitors, and the steady state levels of UCP1 in mature adipocytes may vary depending on physiological and pathological conditions, leading to variability in the levels of UCP1 between superficial and deep neck depots amongst specific individuals (Cypess et al., 2013).

**Figure 2.**
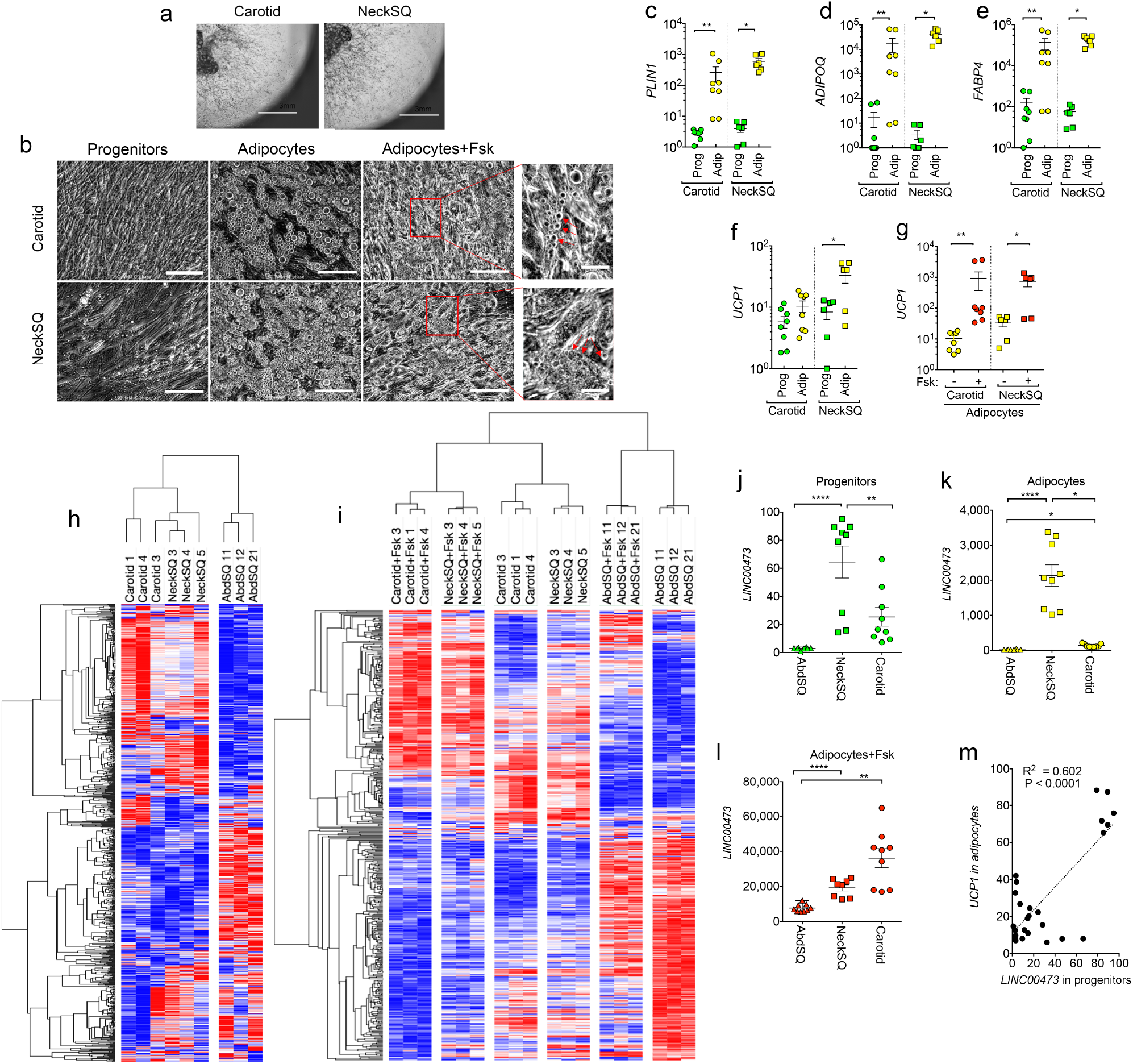
Identification of *LINC00473* as a gene associated with thermogenic depot development. **a.** Example of explants from the indicated depots embedded in Matrigel and cultured for 10 days, showing sprouting and proliferation of progenitors. **b**. Progenitors from Carotid or NeckSQ depots plated on plastic, after differentiation with adipogenic media (adipocytes), and after differentiation and exposure to Fsk daily for the last 5 days in culture (adipocytes+Fsk). Arrowheads in expanded images point to small lipid droplets in cells after Fsk stimulation. Scale bars=200 μm. **c-f** RT-PCR for the genes indicated on the Y-axes, from progenitors or differentiated adipocytes derived from different depots, with or without Fsk stimulation (**g**) as indicated in the x-axes. Values represent the fold-difference over the lowest detected value. Shown are individual values for 2 independent cultures from cells derived from 3 different individuals, and lines are the mean and SEM of these values. Statistical differences were calculated using multiple t-tests, corrected for multiple comparisons using the Holm-Sidak method. *=p<0.05; **p=<0.01. **h**. Unsupervised hierarchical clustering of mean probe intensity values (Affymetrix HTA-2 arrays) of genes in adipocytes derived from Carotid, NeckSQ and AbdSQ depots showing the largest coefficient of variation (range 0.1 to 0.4). i. Unsupervised hierarchical clustering of genes displaying significant differences amongst adipocytes and Fsk-stimulated adipocytes (10 μM Fsk daily for the last 5 days in culture) derived from Carotid, NeckSQ and AbdSQ depots. j-l. RT-PCR of *LINC00473* mRNA in cells derived from the depot indicated in the x-axis. Statistical differences were calculated using the Krustal-Wallis test for non-parametric distributions, corrected for multiple comparisons using the Dunn’s test. *=p<0.05; **p=<0.01; ****p< 0.0001. **m**. Relationship between *LINC00473* values in progenitors and *UCP1* expression in corresponding adipocytes.

To determine whether cells would retain genetic signatures associated with the anatomical origin of their progenitors, we assayed cells from AbdSQ, NeckSQ and Carotid depots using Affymetrix HTA-2 GeneChip microarrays. Unsupervised hierarchical clustering of patterning gene transcripts (*HOX, TBX, MSX, LHX* gene families) from adipocytes differentiated from AbdSQ, NeckSQ and Carotid progenitors resulted in two clearly segregated clusters corresponding to central (AbdSQ) and cranial (NeckSQ and Carotid) depots, and further segregation of the Carotid and NeckSQ depots (Supplementary figure 1). To determine whether adipocytes from different depots would be distinguishable by transcripts other than those associated with patterning genes, we selected the top ~1000 genes that displayed the largest variance (Supplementary table 3). Hierarchical clustering of this set resulted in clear segregation of AbdSQ, NeckSQ and Carotid depots (Figure 2h). This segregation was attributable to variance in genes associated with *PPARγ*, including perilipins, fatty acid binding proteins and adipokines, and to variance in genes associated with epigenetic programing (Supplementary table 4). These results indicate that even after proliferation and differentiation *in vitro*, adipocytes are distinguishable by genetic signatures relevant to adipocyte physiology associated with their depot of origin.

We then asked whether we could find genetic signatures specifically associated with the generation of thermogenic adipocytes in specific depots. For this purpose, we conducted a multi-group differential expression analysis comparing adipocytes from AbdSQ, NeckSQ and Carotid progenitors, with or without Fsk stimulation. Unsupervised hierarchical clustering of genes that were differentially expressed among all conditions (385 genes) resulted in 6 clearly segregated clusters, indicating that the response to Fsk varies as a function of depot of origin (Figure 2i). To determine which genes contributed most to this depot-specific responsiveness, we calculated the median absolute deviation of transcripts in this set. The topmost gene transcript in this analysis corresponded to *LINC00473* (Supplementary Table 5).

The finding of *LINC00473* as the transcript most associated with depot-specific responsiveness to Fsk stimulation, and its strong induction by NE in primary brown adipocytes (Figure 1) suggested it as a candidate in mediating the development of these cells. If so, we hypothesized that *LINC00473* levels would vary as a function of depot of origin at early stages of adipocyte differentiation. To test this hypothesis, we used RT-PCR to determine the levels of *LINC00473* in progenitors from each of the three depots and its response to differentiation (Figure 2j-l). We find that levels of *LINC00473* were higher in progenitors derived from the NeckSQ or Carotid compared to AbdSQ depots (Figure 2j) and increased in response to differentiation in cells from all depots, but to a significantly higher extent in cells from NeckSQ or Carotid compared to AbdSQ (Figure 2k). As expected from the global gene expression analysis, *LINC00473* was strongly induced by Fsk, more so in adipocytes from NeckSQ and Carotid depots compared to those from AbdSQ (Figure 2l). The levels of *LINC00473* in progenitor cells were directly correlated with the levels of *UCP1* seen after differentiation and Fsk stimulation (Figure 2m). These results are consistent with a role for *LINC00473* expression in determining the fate of adipocyte progenitors into the thermogenic phenotype.

To obtain clues onto which biological pathways could be regulated through *LINC00473*, we searched for those genes that were most correlated with its expression levels across all conditions. Enrichment analysis of the 136 genes correlating with *LINC00473* with a Pearson correlation coefficient higher than 0.85 (Supplementary table 6) revealed that the most significantly overrepresented molecular functions were associated with cAMP signaling, including multiple phosphodiesterases and PKA isoforms (Figure 3a, Supplementary Table 7).

We conducted further studies to determine the relationship between *LINC00473* and cAMP signaling in the context of the development and function of thermogenic adipocytes. Four different alternative spliced isoforms of *LINC00473* were found to be expressed, one of which preferentially responded to NE (Figure 3b) and was induced rapidly and reversibly in response to the hormone (Figure 3c). To investigate the relationship between *LINC00473* and cAMP signaling, we analyzed the effects of inhibitors of adenylate cyclase (Figure 3d). Induction of *LINC00473* by NE was dependent on adenylate cyclase activity, placing it downstream of cAMP signaling. Induction of *LINC00473* in response to Fsk preceded the induction of *UCP1*, consistent with a role for *LINC00473* in modulating subsequent *UCP1* expression in response to cAMP (Figure 3e). *LINC00473* induction was also more sensitive to cAMP signaling, responding to lower doses of Fsk compared to *UCP1*, similarly to *NR4A3*, which is an example of a canonically cAMP responsive gene (Figure 3f). Thus, both the time course and sensitivity of responses are consistent with *LINC00473* being induced by, and modulating responses to, cAMP. The functions of long non-coding RNAs are very diverse, ranging from control of gene expression to cytoplasmic scaffolding. To define the potential locus of *LINC00473* function, we compared its subcellular localization in comparison to that of *MALAT1*, a LincRNA known to be abundant and restricted to the nucleus. *MALAT1* was more than 100-fold enriched in the nuclear compared to the cytoplasmic fractions, and did not respond to Fsk stimulation (Figure 3g). In contrast, *LINC00473* was detected both in the nucleus and the cytoplasm in Fsk-stimulated cells (Figure 3h). Similar results were observed in adipocytes derived from SClav in response to NE (not illustrated).

**Figure 3.**
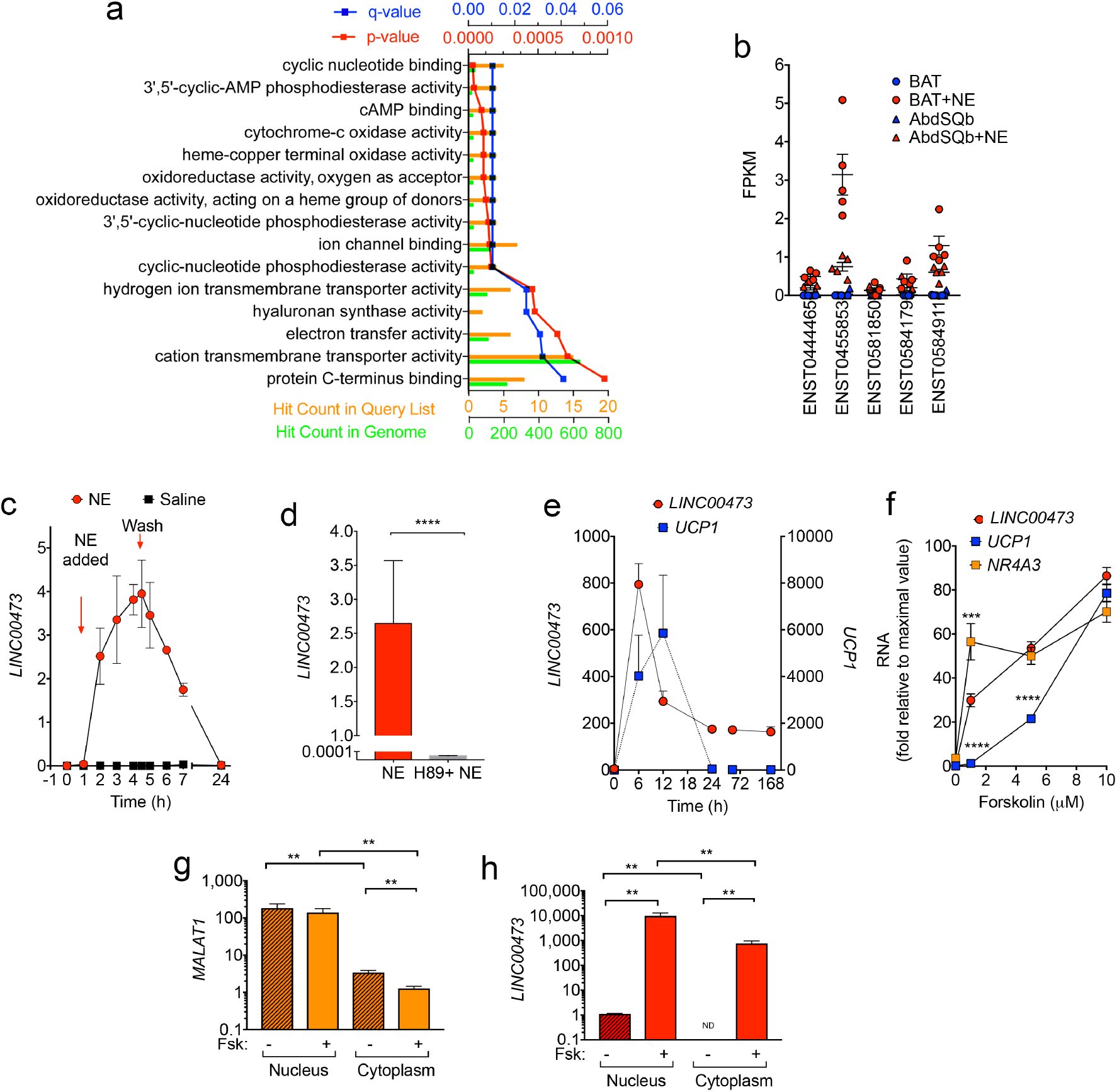
Relationship between *LINC00473* and cAMP signaling. **a**. Pathway enrichment analysis of genes correlated (Pearson correlation coefficient > 0.85, Supplementary Table 5) with *LINC00473* expression in adipocytes derived from Carotid, NeckSQ and AbdSQ depots with and without Fsk stimulation. **b**. Levels of alternative spliced isoforms of *LINC00473* in primary adipocytes from SClav after NE stimulation as in Figure 1. **c**. Levels of *LINC00473* in primary adipocytes from SClav after NE stimulation, in cells treated with vehicle or the adenylate cyclase inhibitor H89. **d**. Kinetics if *LINC00473* induction in primary adipocytes from SClav after NE addition (NE-added) and removal (Wash). **e**. Kinetics of *LINC00473* and *UCP1* induction in adipocytes after addition of 10 μM Fsk at t=0. Values are means and SEM of duplicate cultures from cells derived from 2 different individuals. **f**. Concentration-dependency of *LINC00473, UCP1* and NR4A3 induction in adipocytes after 6h of stimulation with the concentration of Fsk indicated on the x-axis. Statistical differences were calculated using multiple t-tests, corrected for multiple comparisons using the Holm-Sidak method ***p<0.001; ****p< 0.0001. **g-h**. RT-PCR for MALAT1 (**g**) and *LINC00473* (**h**) in the nuclear and cytoplasmic fractions of adipocytes treated without or with 10 μM Fsk for 6 h prior to fractionation. Shown are means and SEM of 5 independent cultures from cells derived from 2 individuals. Statistical differences between pairs were calculated using the Mann-Whitney test, **=P<0.01.

Given the cytoplasmic localization of *LINC00473*, we tested whether siRNA knockdown would interfere with its induction by Fsk. Significant knockdown of *LINC00473* following Fsk-stimulation was seen with two oligonucleotides initially tested (Figure 4a), and subsequent experiments were conducted with a single oligonucleotide that displayed the greatest silencing effect. Transfection of the silencing oligonucleotide resulted in an 80% decrease in *LINC00473* induction following Fsk addition (Figure 4b), and was accompanied by a significant suppression of *UCP1* induction at all time points tested (Figure 4c). The effect of *LINC00473* silencing was less pronounced on the Fsk-induction of *NR4A3* (Figure 4d), suggesting that the actions of *LINC00473* are selective for some cAMP responses. Decreased induction of *UCP1* upon *LINC00473* silencing was seen at all concentrations of Fsk tested (Figure 4e,f), indicating that the effect was due to a loss of responsiveness rather than a decreased sensitivity to the drug. To determine whether impaired induction of *LINC00473* would have a functional effect, we measured oxygen consumption, which is an aggregate of mitochondrial abundance and coupling. Knockdown of *LINC00473* resulted in impaired induction of respiration after addition of NE, which was sustained in the presence of uncoupling, suggesting of an overall decrease in mitochondrial capacity (Figure 4g).

**Figure 4.**
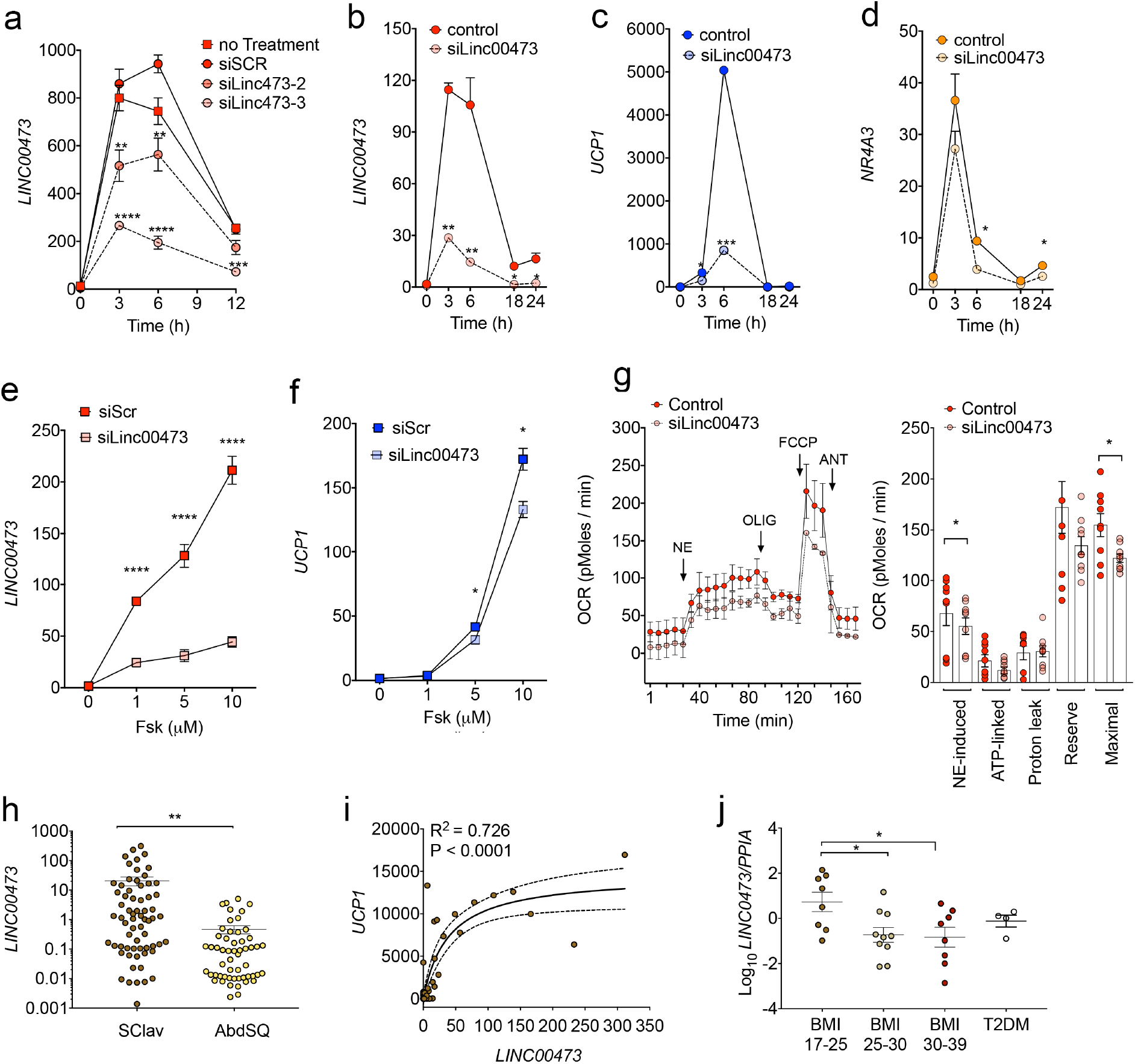
Functional role of *LINC00473.* **a-d**. RT-PCR of genes indicated in the Y-axis after exposure to 10 μM Fsk for the times indicated in the x-axis, from cells treated with indicated oligos 48h prior to extraction. **e-f**. RT-PCR of the genes indicated on the y-axes using RNA extracted after 6h of treatment with the indicated concentrations of Fsk. Values are the mean and SEM of 4 independent cultures of cells from 2 individuals. Statistical differences relative to siScr for **a-f** were calculated using multiple t-tests, corrected for multiple comparisons using the Holm-Sidak method *p<0.05; **p<0.01;***p<0.001; ****p< 0.0001. **g**. Oxygen consumption in primary adipocytes at day 12 of differentiation, treated with scrambled or *LINC00473* targeted siRNAs 72 h prior to assay. Indicated are the times of addition of norepinephrine (NE), Oligomycin (OLIG), FCCP, and rotenone/antimycin (ANT) at concentrations indicated in Materials and Methods. Values are means and SEM of three experiments; mitochondrial respiratory parameters were calculated using three paired time points per condition per experiment, as follows: NE-induced=NE-Basal; ATP-linked=NE-OLIG; Proton leak=ANT-OLIG; Respiratory reserve capacity=FCCP-Basal; Maximal Respiratory Capacity=FCCP-ANT. Bars are the means and SEM. Statistical significance between Control and *siLINC00473* were calculated using paired, 2-tailed student t-tests. *=p<0.05. **h**. Levels of *LINC00473* in tissue sampled from supraclavicular or adominal adipose tissue. Symbols represent FPKM values from RNASeq from each individual, and shown are the means and SEM of the population. **i**. Correlation between *UCP1* and *LINC00473* values in the cohort. **j**. RT-PCR of *LINC00473* in tissue sampled from supraclavicular adipose tissue from individuals with the conditions depicted in the x-axis. Values are the expression of *LINC00473* relative to PP1A used as a housekeeping control. Statistical differences relative to BMI 17-25 were calculated using one-way ANOVA, with Dunnett’s correction for multiple comparisons. *p<0.05

The lack of a homologue of *LINC00473* in mice prevents us from conducting functional experiments in an *in vivo* physiological context. Nevertheless, in humans, obesity and type-2 diabetes have been associated with impairments in thermogenic adipose tissue activity (Saito et al., 2009; Scheele and Nielsen, 2017). To determine whether levels of *LINC00473* could be functionally important for the generation or maintenance of thermogenic adipose tissue in humans, we analyzed its expression in supraclavicular adipose tissue from a large cohort of humans with differing BMI and diabetes status. Levels of *LINC00473* were significantly higher in biopsies from supraclavicular than from matched abdominal subcutaneous adipose tissue (Figure 4h). Moreover, *LINC00473* levels in supraclavicular biopsies correlated strongly with levels of *UCP1* (Figure 4i), and decreased significantly with increasing BMI (Figure 4j). *LINC00473* levels were also lower in subjects with type-2 diabetes (Figure 4j), who are known to display decreased thermogenic adipose tissue. Together with *in vitro* knockdown results, these data suggest that *LINC00473* plays a pivotal role in the generation and maintenance of thermogenic adipose tissue in humans.

## Discussion

Adipose tissue fulfills numerous physiological roles, which are dictated by the properties of adipocytes that compose adipose tissue depots. There are three well-recognized types of adipocytes, which differ fundamentally in biological role. White adipocytes primarily control fuel storage, and brown adipocytes regulate thermogenesis. Beige adipocytes have thermogenic properties but are interspersed within white adipose tissue. Most studies on the mechanisms of development of adipocyte subtypes have come from studies in mice. While there are many similarities between mice and humans, some of the features of mouse adipocytes are not conserved. For example, markers for brown and beige/brite adipocytes identified in mouse do not identify similar adipose types in humans. Therefore, it is important to identify conserved and non-conserved mechanisms involved in adipose tissue development, and to understand the role of depot-dependent function.

Using two different approaches our groups independently identified the same candidate, *LINC00473*, as a determinant of thermogenic fat identity in humans. The combination of our two approaches provides power to identify robust, conserved mechanisms as well as stages in adipocyte development at which these mechanisms operate. In one approach, conventionally cultured pre-adipocytes from the stromal vascular fraction identified *LINC00473* as a gene expressed specifically in adipocytes differentiated from the supraclavicular region, in response to NE stimulation. In the second approach, we generated progenitor cells and differentiated adipocytes from different depots. We found that all depots, Carotid, NeckSQ and AbdSQ, contain progenitors that can give rise to thermogenic fat cells. To find those factors that could determine the higher preponderance of thermogenic adipocytes in the supraclavicular region, we searched for genes that would be differentially expressed between depots, and would be more strongly induced by Fsk in cells from thermogenic compared to non-thermogenic depots. *LINC00473* was the gene that most varied as a function of anatomical localization and depot-dependent sensitivity to stimulation. Besides being enriched in progenitors derived from thermogenic adipose depots, *LINC00473* expression in these cells predicted *UCP1* levels in the differentiated stage. These findings support a role for LINC00473 in the development of thermogenic adipocytes. The further induction of *LINC00473* upon differentiation and after activation by norepinephrine and forskolin supports an additional role as a regulator of thermogenic activity in mature adipocytes.

Long non-coding RNAs are increasingly recognized to control cell development, and several have been identified as required for adipogenesis (Sun et al., 2013). In recent studies, Linc RNAs associated specifically with brown adipocyte generation have also been identified. However, many of these are also required at the differentiation stage. (Li et al., 2016; Liu et al., 2017; Nuermaimaiti et al., 2018; Xiao et al., 2015; Xiong et al., 2018). In contrast, our knockdown studies indicate that *LINC00473* is not required for differentiation, but rather for more specific aspects of thermogenic adipocyte function, as discussed below. Interestingly, *LINC00473* is a primate-specific transcript with no readily identifiable orthologues in other species. This observation is consistent with the reports on a rapid evolution occurring in noncoding sequences with a nucleotide substitution of 90%, compared to a substitution rate of ~10% in protein-coding sequences (Ward et al., 2015). However, RNAs can maintain a conserved secondary structure (Torarinsson et al., 2006), and therefore a functional orthologue of *LINC00473* may exist in mice.

The mechanisms by which *LINC00473* might determine thermogenic adipose tissue identity are likely to be related to modulation of cAMP signaling, which is central to catecholamine-induced thermogenic induction. This possibility is suggested by pathway analysis of genes most closely correlated with *LINC00473* expression across cells and depots, which correspond to cAMP signaling pathways. Moreover, *LINC00473* has been mechanistically associated with cAMP signaling in numerous contexts (Chen et al., 2016; Liang et al., 2016; Pruunsild et al., 2017). *LINC00473* has been reported to interact with NONO, a protein that interacts with cyclic AMP–responsive-element–binding protein (CREB) - regulated transcription coactivator (CRCT), which is essential in CREB transcriptional regulation (Chen et al., 2016). CREB directly and through interaction with NR4A3, (Cannon and Nedergaard, 2004; Kozak et al., 1994; Yubero et al., 1998; Yubero et al., 1994), activates the *UCP1* promoter (Kumar et al., 2008; Myers et al., 2009; Pearen and Muscat, 2010). It is possible that *LINC00473* may coordinate the actions of transcription factors that regulate the *UCP1* promoter.

In addition to a nuclear function, *LINC00473* may operate in the cytoplasm. This is suggested by the large induction of cytoplasmic *LINC00473* upon stimulation by Fsk, which was not observed for our control lncRNA, *MALAT1.* Moreover, its knockdown impaired NE-stimulated and maximal respiration, without affecting proton leak. This suggests that *LINC00473* may play a role in establishing mitochondrial framework necessary for thermogenesis, with roles other than control of *UCP1* expression, possibly at the level of the cytoplasm. The dual localization of *LINC00473* to nucleus and cytoplasm could also relate to a multifunctional role in the establishment and function of thermogenic adipose tissue. In progenitor cells, nuclear *LINC00473* levels may regulate proliferation, and thereby the abundance of thermogenic adipocytes, whereas in the cytoplasm it may regulate responsiveness to cAMP and thereby acute thermogenic activity. Future studies will focus on specific molecular interactions in nucleus and cytoplasm, under basal and stimulated conditions, in cells from different depots. These studies will not only help us understand adipocyte function, but will shed insight on many possible distinct roles of LINC RNAs.

## METHODS

Detailed methods are provided below and include the following:

- KEY RESOURCES TABLE
- CONTACT FOR REAGENT AND RESOURCE SHARING
- EXPERIMENTAL MODEL AND SUBJECT DETAILS

- Human Subjects
- METHOD DETAILS

- Cell Culture
- Bodipy staining
- RNA isolation and reverse transcriptase for primary adipocytes cultures
- RNA-Sequencing.
- RNA extraction of cells derived from human adipose explants
- Affymetrix arrays
- Cell fractionation
- *In vitro* gene silencing of primary adipocytes.
- *In vitro* gene silencing of cells derived from human adipose explants
- Seahorse
- QUANTIFICATION AND STATISTICAL ANALYSIS
- DATA AND SOFTWARE AVAILABILITY

## Supplemental Information

Supplemental information includes 1 figure and 6 tables.

## Author Contributions

SC and SN supervised this work. SN, SC, TF, CS, KVT, TD, SYM, CNB, BKP: hypothesis generation, conceptual design, data analysis, and manuscript preparation. TD, SYM, SC, KVT, CNB, RRR, KM, NJ, HW, NZJ conducting experiments and data analysis.

## Acknowledgements

This study was supported by NIH grant DK089101-04 to SC. We acknowledge the use of services from the UMASS Bioinformatics Core, supported by NIH CTSA grant UL1 TR000161-05, and from the UMASS Genomics Core. KVT is supported by NIH grant 5T32HL120823-03. The Centre for Physical Activity Research (CFAS) is supported by a grant from TrygFonden. During the study period, the Centre of Inflammation and Metabolism (CIM) was supported by a grant from the Danish National Research Foundation (DNRF55). SN was further supported by the Danish Council for Independent Research, Medical Sciences (4092-00492B). NZJ and TJ were funded by Danish Diabetes Academy supported by the Novo Nordisk Foundation. CIM/CFAS is a member of DD2 - the Danish Center for Strategic Research in Type 2 Diabetes (the Danish Council for Strategic Research, grant no. 09-067009 and 09-075724).

## STAR+METHODS KEY RESOURCES TABLE

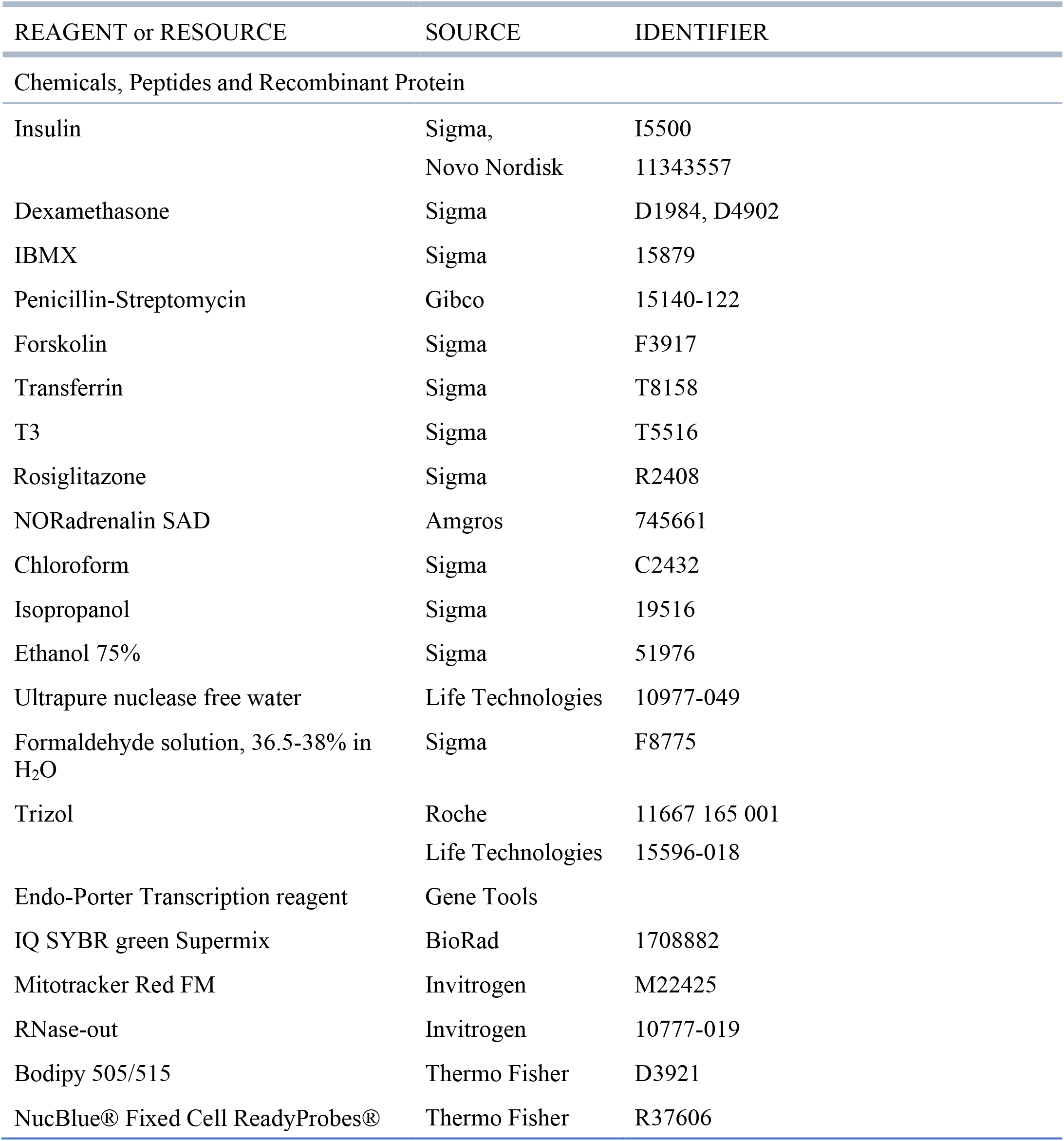
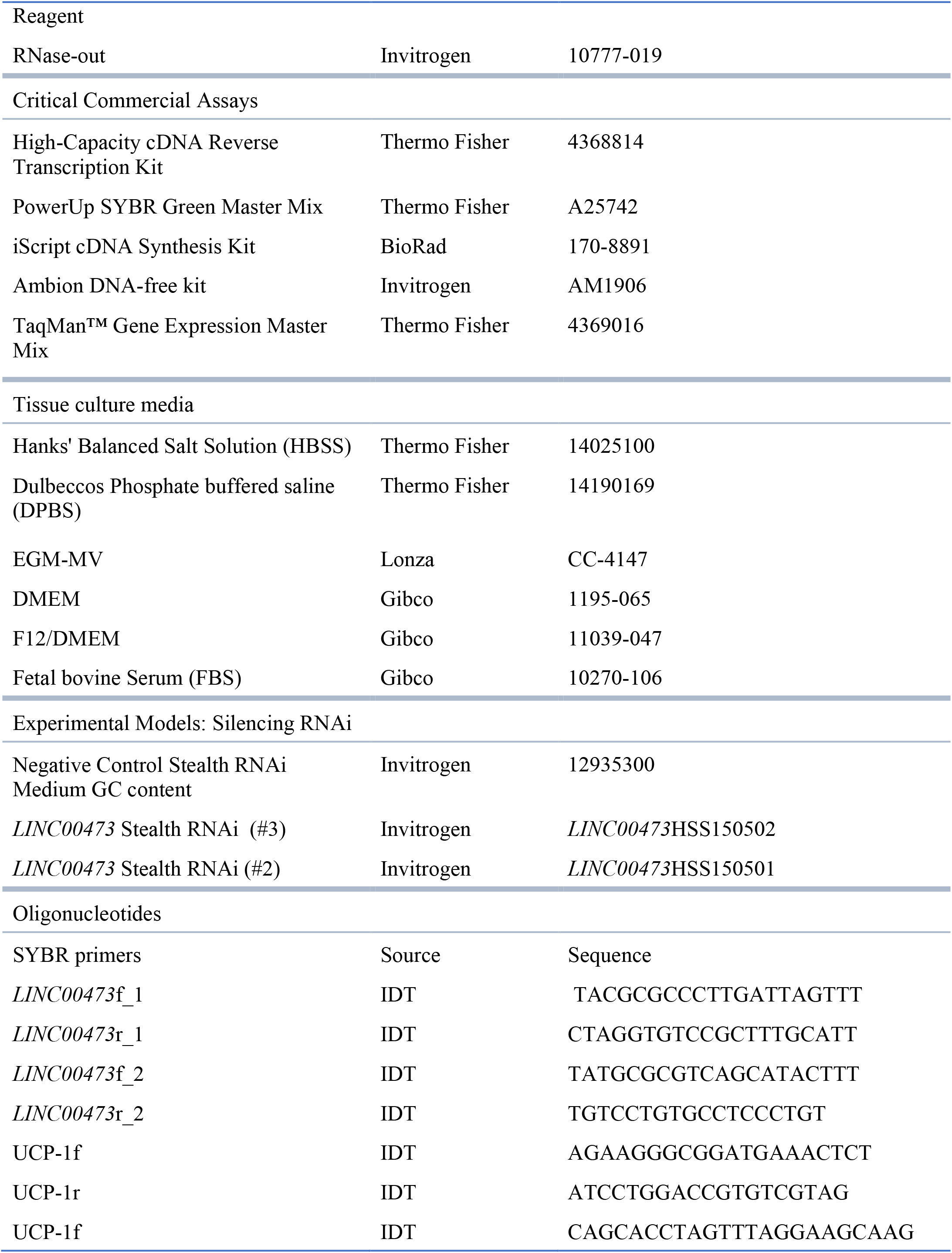
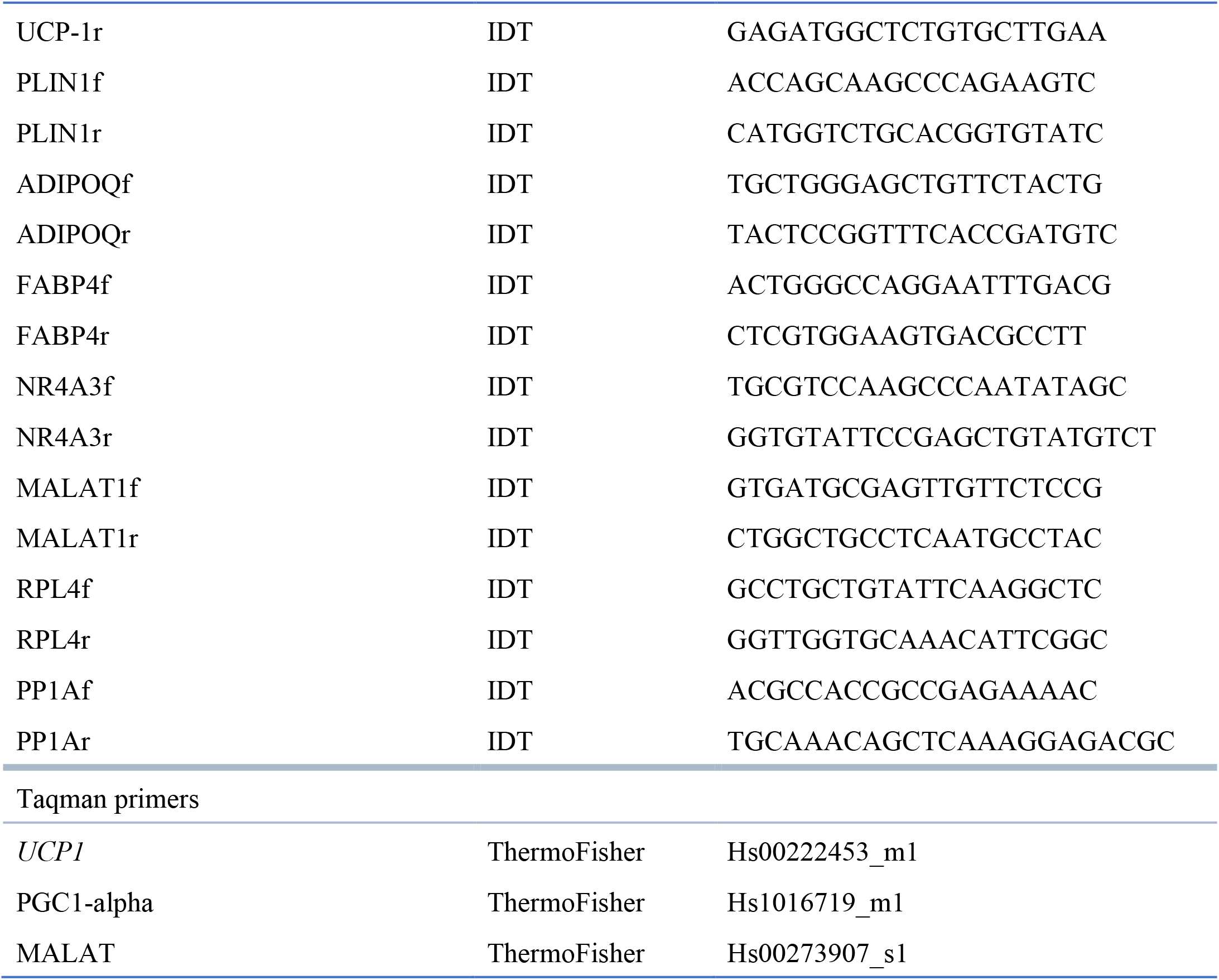

## CONTACT FOR REAGENT AND RESOURCE SHARING

Further information and requests for resources and reagents should be directed to and will be fulfilled by Silvia Corvera

## EXPERIMENTAL MODEL AND SUBJECT DETAILS

### Human Subjects

The study has two cohorts of patients, one of which was based at the UMass Memorial Health Care Center and the other was based at clinics of the Otorhinolaryngology, Head and Neck Surgery and Audiology Departments of Rigshospitalet / Gentofte Hospital and the Department of Otorhinolaryngology, Head, Neck and Maxillofacial Surgery, Zealand University Hospital of Copenhagen, Denmark. All study subjects were age of 23-82 (32% males), not pregnant or incarcerated. The clinical characteristics of the human subjects are listed in Table 1 of Supplementary Information. All participants gave written informed consent. Both studies were performed according to the Declaration of Helsinki and NIH guidelines.

At the UMass Memorial Health Care Center, samples were collected from patients undergoing carotid endarterectomies and panniculectomies. In brief, carotid adipose tissue was collected by removing perivascular fat surrounding the carotid artery during carotid endarterectomies. Neck subcutaneous tissue was collected by sampling the subcutaneous adipose tissue approximately above the area where perivascular tissue was taken. Abdominal adipose tissue was collected from discarded tissues of patients undergoing panniculectomies. Tissue was harvested, placed in EGM-2 MV. Adipose tissue was then minced in 1mm pieces and embedded in Matrigel as detailed below. This study was approved by the University of Massachusetts Institutional Review Board, IRB H00001329.

At the Gentofte Hospital and the Zealand University Hospital of Copenhagen (Denmark), the human cohort is well-characterized and described in detail in separate manuscript (Jespersen et al 2018, under preparation). In brief, 35 patients scheduled for surgery due to benign goiter were included in the main study, and 36 were additionally included in the biopsy part only. The material was collected from 2014 and October 2016 at the outpatient clinics of the Otorhinolaryngology, Head and Neck Surgery and Audiology Departments of Rigshospitalet / Gentofte Hospital and the Department of Otorhinolaryngology, Head, Neck and Maxillofacial Surgery, Zealand University Hospital of Copenhagen, Denmark. Apart from thyroid malignancy and inability to provide informed consent, there were no specific exclusion criterions. The Scientific-Ethics Committees of the Capital Region of Denmark approved the study protocol (HD-2009-020). The characteristics listed in Table 1 includes the mean BMI, gender and age for the subjects included in the adipose tissue depot comparison presented in figure 4H. A subgroup of 30 individuals were further examined for blood glucose levels after an Oral Glucose Tolerance Test (OGTT). One subject was excluded from the analysis due to high thyroid hormone levels. The characteristics for the remaining 29 subjects are presented SI Table 1, last section. The subjects were divided into three BMI groups, normal weight (BMI < 25), overweight (BMI 25-30) and obese (BMI > 30) and a Type 2 diabetes group (T2DM). The four individuals in the T2DM group were classified based on the blood glucose levels in the OGTT (definition from the World Health Organization (WHO)). A detailed description of the OGGT procedure is described in detail in (Jespersen et al 2018, under preparation). The subcutaneous abdominal biopsies were obtained using a modified Bergström needle biopsy procedure as previously described after induction of general anesthesia and immediately prior to initiation of surgery. The supraclavicular biopsies were obtained by the surgeon from the deep neck fatty tissue depots (Bergstrom, 1975). Biopsy samples were homogenized in TRIzol (Invitrogen, Carlsbad, CA, USA) using a Tissuelyser (Qiagen, Valencia, CA, USA) and total RNA was then isolated as described in the method section “RNA isolation and reverse transcriptase for primary adipocytes cultures”.

## METHOD DETAILS

### Cell culture

#### Cell culture of human primary adipocytes from SVF

As previously described, preadipocytes were isolated from supraclavicular and abdominal subcutaneous adipose tissue biopsies (Jespersen et al., 2013). Biopsies were minced and digested in DMEM/F12 (containing collagenase II (1 mg/ml)(Sigma Aldrich) and fatty acid-free bovine serum albumin (15 mg/ml) (Sigma Aldrich) for 20 min at 37 °C during gentle shaking. Following digestion, the suspension was filtered through a 70-micron cell strainer and left to settle for 5 min. The liquid phase below the upper lipid phase was aspirated using a syringe and passed through a 30-micron filter. The cell suspension was spun down at 800 g for 7 min and washed with DMEM/F12. Preadipocytes were resuspended in DMEM/F12, 1% penicillin/streptomycin, 10% fetal bovine serum (FBS) (Life technologies) and seeded in a 5-ml culture flask. Media was changed the day after isolation and then every second day until cells were 80% confluent and then split into a 10-cm dish (passage 1). For the cell experiments, preadipocytes were cultured in 100 mm and 6 cm culture dishes containing DMEM/F12, 10% FBS, 1% Penicillin-Streptomycin (all from Invitrogen) and 1 nM Fibroblast growth factor-acidic (FGF-1) (ImmunoTools). The cells were grown at 37°C in an atmosphere of 5% CO2 and the medium was changed every second day. Adipocyte differentiation was induced two days after preadipocyte cultures were 100% confluent by treating cells with DMEM/F12 containing 1% Penicillin-Streptomycin, 0.1 μM dexamethasone (Sigma-Aldrich), 100 nM insulin (Actrapid, Novo Nordisk or Humulin, Eli Lilly), 200 nM rosiglitazone (Sigma-Aldrich), 540 μM isobutylmethylxanthine (IBMX) (Sigma-Aldrich), 2 nM T3 (Sigma-Aldrich) and 10 μg/ml transferrin (Sigma-Aldrich). After three days of differentiation, IBMX was removed from the cell culture media. The cell cultures were left to differentiate for an additional nine days, with medium change third day. Following 12 days of differentiation, cells were harvested for RNA, protein. When stated in the figure legend, cells were stimulated with 10 μM norepinephrine (Sigma-Aldrich) for 4 hr before RNA and protein were isolated. Two hours prior to the norepinephrine stimulation, old medium was replaced by DMEM/F12 (Life technologies) containing 1% penicillin-streptomycin.

#### Cell culture of progenitor cells derived from human adipose explants

We collected carotid peri-vascular and neck subcutaneous adipose tissues from carotid endarterectomies with no a-priori selection of individual donors. The characteristics of patients from which tissues were used for indicated experiments are described in Supplementary Table 1. Detailed methods for culture adipose tissue explants and harvesting of single cells from explant growth are published (Rojas-Rodriguez et al., 2014). In brief, cell suspensions from capillary growth were obtained using dispase, and plated on standard tissue culture plates. Growth and passaging of these cells was done using EGM-2 MV. To induce adipogenesis we used a minimal adipogenic cocktail of DMEM +10% FBS, 0.5 mM 3-isobutyl-1-methylxanthine, 1μM dexamethasone, and 1μg/ml insulin (MDI) for 72h. The medium was then replaced with DMEM plus 10% FBS. Subsequently, 50% of the medium was replaced with fresh medium every other day. Adipocyte markers were measured by RT-PCR in RNA extracted from 3 explants per condition.

### Bodipy staining

Fully differentiated adipocytes were fixed in 4% Formaldehyde for 15 min and washed with DPBS (Gibco) three times. Bodipy (Thermo Fisher) was diluted in HBSS to a final concentration at 0.5mM and incubated with fixed cells for 20 min. After washing with DPBS, 1 drop NucBlue™ Fixed Cell ReadyProbes™ Reagent (Thermo Fisher) pr. ml HBSS was added for staining of the nucleus (8 min incubation). Pictures of cells were taken with EVOS Auto microscope (Thermo Fisher).

### RNA isolation and reverse transcriptase for primary adipocytes cultures

Total RNA (200 ng) was reverse-transcribed using cDNA high capacity kit (Applied Biosystems) according to the manufacturer’s protocol. cDNA samples were loaded in triplicates and qPCR was performed using ViiA7 Sequence Detection system (Applied Biosystems, Foster City, CA, USA) according to the manufacturer’s protocol using either PowerUp SYBR Green Master Mix (Thermo Fisher) or TaqMan™ Universal PCR Master Mix (Thermo Fisher). SYBR based primers were designed using Roche Applied Science Assay Design Center (Roche) and checked for specificity using Primer-Blast (NCBI)

### RNA-Sequencing

RNA (1000 ng) was extracted from adipocytes using the Trizol method. RNA sequencing was performed by BGI (Hong Kong) using 1000 ng RNA for the TruSeq cDNA library construction (Illumina). 3Gb data was generated pr sample on a HiSeq 2000 sequencer (Illumina). A 91-paired end sequencing strategy was used for the project. Overall read quality was assessed using FastQC http://www.bioinformatics.babraham.ac.uk and the following pre-processing steps where performed using the Fastx toolkit (http://hannonlab.cshl.edu) and PRINSEQ: 7 nt were clipped off from the 5’ end of every read (Schmieder and Edwards, 2011). The reads were then filtered to remove all N-reads. The 3’ ends were then trimmed to and the the reads filtered to minimum Q25 and 50 bp length. Reads were then mapped with tophat2 to the human genome GRCh38 Ensembl release 77 (Kim et al., 2013). Read counts where imported into R, and DESeq2 was used for identifying differential expression (Love et al., 2014). For the isoform analysis, fpkm values from cufflinks were used (Roberts et al., 2011; Trapnell et al., 2010).

### RNA extraction of cells derived from human adipose explants

Media was aspirated from the well and cells were washed 2X with PBS. TriPure Trizol reagent was added to the cells and incubated at room temperature for 5 minutes. Cells were collected into a GentleMACS M tube and dissociated using the GentleMACS Dissociator (Miltenyi Bio) Program RNA01.01. Tubes were centrifuged for 3 minutes at 800RPM and the mixture was transferred to a 2ml collection tube. Chlorofom was added in a 1:5 ratio to the tripure/cell mix and tubes were inverted to mix, then incubated at room temperature for 3-5 minutes. Aqueous phase separation was performed and the RNA-containing layer was mixed with an equal volume of 100% Isopropanol and incubated overnight at −20 degrees for precipitation. RNA was pelleted and washed with 80% ETOH, and eluted in nuclease-free water. Nucleotide concentrations were determined using Nanodrop 2000. 1μg of RNA was reverse transcribed using iScript cDNA Synthesis Kit (BioRad). Primer sequences are shown in supplemental Table 6.

### Affymetrix arrays

Total RNA was isolated using TRIzol as above. Affymetrix protocols were followed for the preparation of cRNA, which was hybridized to HTA-2.0 arrays. Raw expression data collected from an Affymetrix HP GeneArrayScanner was normalized across all data sets using the RMA algorithm as implemented by the Affymetrix Expression Console. Expression analysis was performed using the Affymetrix Transcriptome Analysis Console v.3.0.

### Cell fractionation

Human preadipocytes were seeded at a density of 3×10^6 cells per plate into three 10cm plates per condition, and grown to confluence for 72h. Plates were differentiated with MDI media for a total of 7 days (see differentiation protocol). Six hours prior to collection one half of the plates were stimulated with 10uM Forskolin (Sigma, F3917). Cells were collected by trypanization and washed 1x with PBS. Cells were pelleted and stored at −80 degrees overnight. Cells were then re-suspended into 2ml of cracking buffer (50mM Hepes pH 7.9, 3mM MgCl2, 1mM DTT, 0.25M Sucrose, 40U/ml RNase-out). Samples were loaded into a 3ml syringe and passed through a cold Balch homogenizer twenty times. Collected supernatant was centrifuged at 700xg for 10 minutes at 4 degrees. Supernatant was collected and stored for analysis of the cytoplasmic fraction. Pellets were washed one time with buffer and spun again at 700Xg for 10min at 4 degrees. Supernatant was carefully removed from remaining pellet and the pellet was washed once with 1X PBS and pelleted for RNA extraction. RNA was extracted from the cytoplasm and nuclear fractions using Trizol-chloroform extraction method (TriPure, Roche). Eluted RNA was treated with DNase following extraction (Invitogen).

### *In vitro* gene silencing of primary adipocytes

Transfections on adipocytes were performed using siRNA pools consisting of four siRNA oligos specifically targeting four different sites of target *LINC00473* (On Target Plus, R-032718-00-0005, Dharmacon). Transfections were performed using 10 ul Lipofectamine^®^ RNAiMAX Transfection Reagent (Thermo Fisher Scientific) with 20 nm nM of siRNA, at day 11 of differentiation in antibiotic-free cell culture media (Opti-MEM^®^, F12/DMEM) for 24h. A scrambled non-specific oligonucleotide (siRNA Scr) was used as control.

### *In vitro* gene silencing of cells derived from human adipose explants

Cells were plated at 1×10^5 cells per well in 24-well dish. After reaching confluency (~3 days), cells were exposed to MDI for 72h, followed by 2 days with DMEM+FBS media alone. Cells were then transfected (day 5 of differentiation) with *LINC00473* or negative control stealth RNAi (Invitrogen) using Endo-Porter Transfection reagent (Gene Tools). 48h later (day 7 of differentiation) cells were stimulated with vehicle or Fsk and 6h later RNA was extracted. For transfection, RNAi and endoporter were combined in a 1.5 eppendorf tube and allowed to complex for 15 minutes at room temperature. DMEM culture media was added to the mixture to reach a final concentration of 500nM siRNA and 7.5 μM Endo-Porter reagent. All of the media on the cells was replaced with either siLINC00473/Endo-Porter or siSCR/Endo-Porter-containing media and cells were incubated for 48h at 37 degrees and 5% CO2. 3-24 hours prior to collection cells were stimulated in DMEM culture media containing 1-10μM Forskolin (Sigma).

### Oxygen consumption

Oxygen consumption was measured using a Seahorse Bioscience XF96 Extracellular Flux Analyser according to the manufacturer’s protocol. Adipocytes were grown until reaching 100% confluency and were then seeded in seahorse plates at a 1:1 ratio, and differentiated as described above. Experiments were performed on day 12 of differentiation on cells in passage three and Knock down experiments were performed as described above. Oxygen consumption rate was assessed in 4 primary brown adipocyte cultures. The results were extracted from the Seahorse Program Wave 2.2.0. Baseline measurements of OCR were performed for 30 minutes before NE or saline was added and measurements of the concomitant responses were recorded for 60 minutes. All other states were induced using the seahorse XF cell mito stress test kit according to the manufactures protocol. After 90 minutes, leak state was induced by adding Oligomycin, which inhibits the ATP synthase. Leak state measurements were performed for 20 minutes, then the ionophore (carbonyl cyanide-4-(trifluoromethoxy) phenylhydrazone) (FCCP), which collapses the proton gradient across the mitochondrial inner membrane resulting in a completely uncoupled state. After an additional 20 minutes Antimycin A and Rotenone were added to inhibit complexes III and I respectively, resulting in only non-mitochondrial respiration. For data analyses OCR was corrected for non-mitochondrial respiration as assessed by the Seahorse XF cell mitochondrial stress test kit Wells were excluded from the data analyses if OCR were +/-20% of the mean in that series of replicate values.

## QUANTIFICATION AND STATISTICAL ANALYSIS

GraphPad Prism 7.0 was used for all analyses. Parametric or non-parametric test were chosen based on results from the D’Agostino-Pearson omnibus normality test, and are described in each figure. Heatmaps were plotted using Morpheus (Broad Institute).

---

**Supplementary Figure 1.**
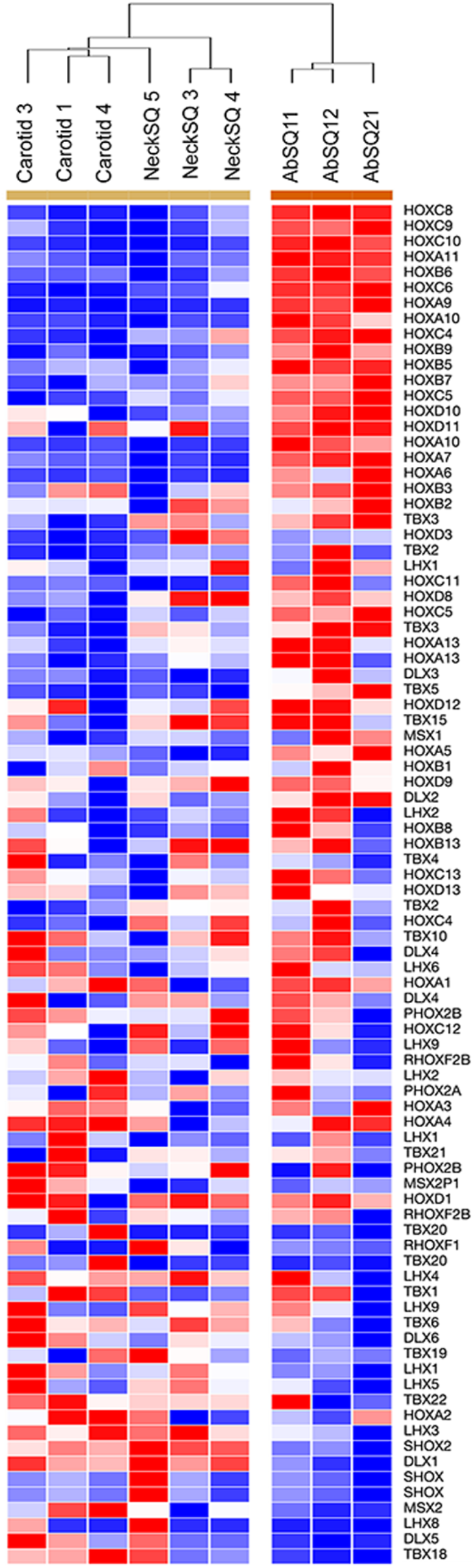
Hierarchical clustering (Spearman, single; rows arranged by nearest neighbor of gene showing highest variance (HOXC8)) of mean probe intensity values for developmental genes, obtained in HTA-2.0 Affymetrix arrays of RNA from adipocytes derived from Carotid, NeckSQ and AbdSQ depots.

Supplementary Table 1: Patient Characteristics

Supplementary Table 2. Genes differentially expressed in response to norepinephrine

Supplementary Table 3: Differentiated adipocyte genes with high variance between depots

Supplementary Table 4: Pathway and enrichment analysis of genes displaying the highest coefficient of variation

Supplementary Table 5: Multi-group differential expression analysis of adipocytes from AbdSQ, NeckSQ and Carotid progenitors with or without Fsk stimulation

Supplementary Table 6: Genes correlated with LINC00473

Supplementary Table 7. Pathway enrichment analysis of genes correlated with LINC00473

*Please note, supplementary tables are placed in separate file due to size.

## REFERENCES

Alvarez-Dominguez, J.R., Bai, Z., Xu, D., Yuan, B., Lo, K.A., Yoon, M.J., Lim, Y.C., Knoll, M., Slavov, N., Chen, S., et al. (2015). De Novo Reconstruction of Adipose Tissue Transcriptomes Reveals Long Non-coding RNA Regulators of Brown Adipocyte Development. Cell Metab 21, 764–776.

Atianand, M., Caffrey, D., and Fitzgerald, K. (2017). Immunobiology of Long Noncoding RNAs. Annu Rev Immunol. 35.

Bai, Z., Chai, X.R., Yoon, M.J., Kim, H.J., Lo, K.A., Zhang, Z.C., Xu, D., Siang, D.T.C., Walet, A.C.E., Xu, S.H., et al. (2017). Dynamic transcriptome changes during adipose tissue energy expenditure reveal critical roles for long noncoding RNA regulators. PLoS Biol 15, e2002176.

Bergstrom, J. (1975). Percutaneous needle biopsy of skeletal muscle in physiological and clinical research. Scand J Clin Lab Invest 35, 609–616.

Berry, R., and Rodeheffer, M.S. (2013). Characterization of the adipocyte cellular lineage in vivo. Nat Cell Biol 15, 302–308.

Cannon, B., and Nedergaard, J. (2004). Brown adipose tissue: function and physiological significance. Physiol Rev 84, 277–359.

Chen, Z., Li, J., Lin, S., Cao, C., Gimbrone, N., and Yang R, F.D., Carper MB, Haura EB, Schabath MB, Lu J, Amelio AL, Cress WD, Kaye FJ, Wu L. (2016). cAMP/CREB-regulated LINC00473 marks LKB1-inactivated lung cancer and mediates tumor growth. J Clin Invest 126, 2267–2279.

Cinti, S. (2001). The adipose organ: morphological perspectives of adipose tissues. Proc Nutr Soc 60, 319–328.

Crewe, C., An, Y.A., and Scherer, P.E. (2017). The ominous triad of adipose tissue dysfunction: inflammation, fibrosis, and impaired angiogenesis. J Clin Invest 127, 74–82.

Cypess, A.M., White, A.P., Vernochet, C., Schulz, T.J., Xue, R., Sass, C.A., Huang, T.L., Roberts-Toler, C., Weiner, L.S., Sze, C., et al. (2013). Anatomical localization, gene expression profiling and functional characterization of adult human neck brown fat. Nat Med 19, 635–639.

Faghihi, M., Modarresi, F., Khalil, A., Wood, D., Sahagan, B., Morgan, T., Finch, C., St Laurent, G.r., Kenny, P., and Wahlestedt, C. (2008). Expression of a noncoding RNA is elevated in Alzheimer’s disease and drives rapid feed-forward regulation of beta-secretase. Nat Med 14 723–730.

Freedman, J., and Miano, J. (2016). Challenges and Opportunities in Linking Long Noncoding RNAs to Cardiovascular, Lung, and Blood Diseases. Arterioscler Thromb Vasc Biol. 2017 Jan;37(1):21–25. Arterioscler Thromb Vasc Biol., 1.

Gupta, R., Shah, N., Wang, K., Kim, J., Horlings, H., Wong, D., Tsai, M., Hung, T., Argani, P., Rinn, J., et al. (2010). Long non-coding RNA HOTAIR reprograms chromatin state to promote cancer metastasis. Nature 464, 1071–1076.

Gutschner, T., and Diederichs, S. (2012). The hallmarks of cancer: a long non-coding RNA point of view. RNA Biol. 9, 703–719.

Harms, M., and Seale, P. (2013). Brown and beige fat: development, function and therapeutic potential. Nat Med 19, 1252–1263.

Hezroni, H., Koppstein, D., Schwartz, M.G., Avrutin, A., Bartel, D.P., and Ulitsky, I. (2015). Principles of long noncoding RNA evolution derived from direct comparison of transcriptomes in 17 species. Cell Rep 11, 1110–1122.

Jeffery, E., Wing, A., Holtrup, B., Sebo, Z., Kaplan, J.L., Saavedra-Pena, R., Church, C.D., Colman, L., Berry, R., and Rodeheffer, M.S. (2016). The Adipose Tissue Microenvironment Regulates Depot-Specific Adipogenesis in Obesity. Cell Metab 24, 142–150.

Jespersen, N.Z., Larsen, T.J., Peijs, L., Daugaard, S., Homoe, P., Loft, A., de Jong, J., Mathur, N., Cannon, B., Nedergaard, J., et al. (2013). A classical brown adipose tissue mRNA signature partly overlaps with brite in the supraclavicular region of adult humans. Cell Metab 17, 798–805.

Kim, D., Pertea, G., Trapnell, C., Pimentel, H., Kelley, R., and Salzberg, S.L. (2013). TopHat2: accurate alignment of transcriptomes in the presence of insertions, deletions and gene fusions. Genome Biol 14, R36.

Kozak, U., Kopecky, J., Teisinger, J., Enerbäck, S., Boyer, B., and Kozak, L. (1994). An upstream enhancer regulating brown-fat-specific expression of the mitochondrial uncoupling protein gene. Mol Cell Biol 14, 59–67.

Kumar, N., Liu, D., Wang, H., Robidoux, J., and Collins, S. (2008). Orphan nuclear receptor NOR-1 enhances 3′,5′-cyclic adenosine 5′-monophosphate-dependent uncoupling protein-1 gene transcription. Mol Endocrinol. 22 1057–1064.

Li, M., Sun, X., Cai, H., Sun, Y., Plath, M., Li, C., Lan, X., Lei, C., Lin, F., Bai, Y., et al. (2016). Long non-coding RNA ADNCR suppresses adipogenic differentiation by targeting miR-204. Biochim Biophys Acta 1859, 871–882.

Liang, X.H., Deng, W.B., Liu, Y.F., Liang, Y.X., Fan, Z.M., Gu, X.W., Liu, J.L., Sha, A.G., Diao, H.L., and Yang, Z.M. (2016). Non-coding RNA LINC00473 mediates decidualization of human endometrial stromal cells in response to cAMP signaling. Sci Rep 6, 22744.

Liu, W., Ma, C., Yang, B., Yin, C., Zhang, B., and Xiao, Y. (2017). LncRNA Gm15290 sponges miR-27b to promote PPARgamma-induced fat deposition and contribute to body weight gain in mice. Biochem Biophys Res Commun 493, 1168–1175.

Love, M.I., Huber, W., and Anders, S. (2014). Moderated estimation of fold change and dispersion for RNA-seq data with DESeq2. Genome Biol 15, 550.

M, H., M, G., D, F., M, G., MJ, K., D, K.-B., AM, K., O, Z., I, A., M, R., et al. (2010). A large intergenic noncoding RNA induced by p53 mediates global gene repression in the p53 response. Cell 142, 409–419.

Mercer, T., Dinger, M., and Mattick, J. (2009). Long non-coding RNAs: insights into functions. Nat Rev Genet. 10, 155–159.

Min, S.Y., Kady, J., Nam, M., Rojas-Rodriguez, R., Berkenwald, A., Kim, J.H., Noh, H.L., Kim, J.K., Cooper, M.P., Fitzgibbons, T., et al. (2016). Human ‘brite/beige’ adipocytes develop from capillary networks, and their implantation improves metabolic homeostasis in mice. Nat Med 22, 312–318.

Mota de Sa, P., Richard, A.J., Hang, H., and Stephens, J.M. (2017). Transcriptional Regulation of Adipogenesis. Compr Physiol 7, 635–674.

Myers, S., Eriksson, N., Burow, R., Wang, S., and Muscat, G. (2009). Beta-adrenergic signaling regulates NR4A nuclear receptor and metabolic gene expression in multiple tissues. Mol Cell Endocrinol 309, 101–108.

Nedergaard, J., and Cannon, B. (2010). The changed metabolic world with human brown adipose tissue: therapeutic visions. Cell Metab 11, 268–272.

Nuermaimaiti, N., Liu, J., Liang, X., Jiao, Y., Zhang, D., Liu, L., Meng, X., and Guan, Y. (2018). Effect of lncRNA HOXA11-AS1 on adipocyte differentiation in human adipose-derived stem cells. Biochem Biophys Res Commun 495, 1878–1884.

Pearen, M., and Muscat, G. (2010). Minireview: Nuclear hormone receptor 4A signaling: implications for metabolic disease. Mol Endocrinol. 24, 1891–1903.

Pruunsild, P., Bengtson, C.P., and Bading, H. (2017). Networks of Cultured iPSC-Derived Neurons Reveal the Human Synaptic Activity-Regulated Adaptive Gene Program. Cell Rep 18, 122–135.

Roberts, A., Trapnell, C., Donaghey, J., Rinn, J.L., and Pachter, L. (2011). Improving RNA-Seq expression estimates by correcting for fragment bias. Genome Biol 12, R22.

Rojas-Rodriguez, R., Gealekman, O., Kruse, M.E., Rosenthal, B., Rao, K., Min, S., Bellve, K.D., Lifshitz, L.M., and Corvera, S. (2014). Adipose tissue angiogenesis assay. Methods Enzymol 537, 75–91.

Rosen, E.D., and Spiegelman, B.M. (2006). Adipocytes as regulators of energy balance and glucose homeostasis. Nature 444, 847–853.

Saito, M., Okamatsu-Ogura, Y., Matsushita, M., Watanabe, K., Yoneshiro, T., Nio-Kobayashi, J., Iwanaga, T., Miyagawa, M., Kameya, T., Nakada, K., et al. (2009). High incidence of metabolically active brown adipose tissue in healthy adult humans: effects of cold exposure and adiposity. Diabetes 58, 1526–1531.

Sanchez-Gurmaches, J., Hung, C.M., and Guertin, D.A. (2016). Emerging Complexities in Adipocyte Origins and Identity. Trends Cell Biol 26, 313–326.

Scheele, C., and Nielsen, S. (2017). Metabolic regulation and the anti-obesity perspectives of human brown fat. Redox Biol 12, 770–775.

Schmieder, R., and Edwards, R. (2011). Quality control and preprocessing of metagenomic datasets. Bioinformatics 27, 863–864.

Simon, M., Wang, C., Kharchenko, P., West, J., Chapman, B., Alekseyenko, A., Borowsky, M., Kuroda, M., and Kingston, R. (2011). The genomic binding sites of a noncoding RNA. Proc Natl Acad Sci U S A. 108, 20497–20502.

Sun, L., Goff, L.A., Trapnell, C., Alexander, R., Lo, K.A., Hacisuleyman, E., Sauvageau, M., Tazon-Vega, B., Kelley, D.R., Hendrickson, D.G., et al. (2013). Long noncoding RNAs regulate adipogenesis. Proc Natl Acad Sci U S A 110, 3387–3392.

Torarinsson, E., Sawera, M., Havgaard, J.H., Fredholm, M., and Gorodkin, J. (2006). Thousands of corresponding human and mouse genomic regions unalignable in primary sequence contain common RNA structure. Genome Res 16, 885–889.

Tran, K.V., Gealekman, O., Frontini, A., Zingaretti, M.C., Morroni, M., Giordano, A., Smorlesi, A., Perugini, J., De Matteis, R., Sbarbati, A., et al. (2012). The vascular endothelium of the adipose tissue gives rise to both white and brown fat cells. Cell Metab 15, 222–229.

Trapnell, C., Williams, B.A., Pertea, G., Mortazavi, A., Kwan, G., van Baren, M.J., Salzberg, S.L., Wold, B.J., and Pachter, L. (2010). Transcript assembly and quantification by RNA-Seq reveals unannotated transcripts and isoform switching during cell differentiation. Nat Biotechnol 28, 511–515.

Tsai, M., Manor, O., Wan, Y., Mosammaparast, N., Wang, J., Lan, F., Shi, Y., Segal, E., and Chang, H. (2010). Long noncoding RNA as modular scaffold of histone modification complexes. Science 329, 689–693.

Virtanen, K.A., Lidell, M.E., Orava, J., Heglind, M., Westergren, R., Niemi, T., Taittonen, M., Laine, J., Savisto, N.J., Enerback, S., et al. (2009). Functional brown adipose tissue in healthy adults. N Engl J Med 360, 1518–1525.

Wang, X., Arai, S., Song, X., Reichart, D., Du, K., Pascual, G., Tempst, P., Rosenfeld, M., Glass, C., and Kurokawa, R. (2008). Induced ncRNAs allosterically modify RNA-binding proteins in cis to inhibit transcription. Nature 454, 126–130.

Ward, M., McEwan, C., Mills, J.D., and Janitz, M. (2015). Conservation and tissue-specific transcription patterns of long noncoding RNAs. J Hum Transcr 1, 2–9.

Xiao, T., Liu, L., Li, H., Sun, Y., Luo, H., Li, T., Wang, S., Dalton, S., Zhao, R.C., and Chen, R. (2015). Long Noncoding RNA ADINR Regulates Adipogenesis by Transcriptionally Activating C/EBPalpha. Stem Cell Reports 5, 856–865.

Xiong, Y., Yue, F., Jia, Z., Gao, Y., Jin, W., Hu, K., Zhang, Y., Zhu, D., Yang, G., and Kuang, S. (2018). A novel brown adipocyte-enriched long non-coding RNA that is required for brown adipocyte differentiation and sufficient to drive thermogenic gene program in white adipocytes. Biochim Biophys Acta 1863, 409–419.

Yubero, P., Barberá, M., Alvarez, R., Viñas, O., Mampel, T., Iglesias, R., Villarroya, F., and Giralt, M. (1998). Dominant negative regulation by c-Jun of transcription of the uncoupling protein-1 gene through a proximal cAMP-regulatory element: a mechanism for repressing basal and norepinephrine-induced expression of the gene before brown adipocyte differentiation. Mol Endocrinol 12 1023–1037.

Yubero, P., Manchado, C., Cassard-Doulcier, A., Mampel, T., Viñas, O., Iglesias, R., Giralt, M., and Villarroya, F. (1994). CAAT/enhancer binding proteins and are transcriptional activators of the brown fat uncoupling protein gene promoter. Biochem Biophys Res Commun. 1994 Jan 28;198(2):653-9 198, 653-659.

Zhao, X.Y., Li, S., Wang, G.X., Yu, Q., and Lin, J.D. (2014). A long noncoding RNA transcriptional regulatory circuit drives thermogenic adipocyte differentiation. Mol Cell 55, 372–382.

